# Patterned membrane recycling balances apical and lateral cell surfaces to confine tissue deformation under stretch

**DOI:** 10.64898/2026.10.20.760495

**Authors:** Elise Trubuil, Jone Isasti-Sanchez, Lenka Backová, Ignacio Arganda-Carreras, Jérôme Solon

**Author notes:** These authors contributed equally to this work (Elise Trubuil, Jone Isasti-Sanchez).

## Abstract

During development, epithelia undergo large stretches, but how tissues confine the resulting deformations and stresses to specific regions is poorly understood. Here, we show that during *Drosophila* dorsal closure, the epidermis confines deformation through a developmentally controlled, patterned increase in apical cell surface. Dorsal cells respond to stretch by expanding their apical surface and flattening, whereas ventral cells do not expand nor flatten, even when over-stretched and despite reorganizing their actomyosin. Both shape changes require Rab11-dependent membrane recycling, showing that cell height and apical area are actively controlled rather than passively set by stretch. The Dpp/Brinker pathway restricts this remodeling to the dorsal epidermis. A vertex model shows that localized remodeling is sufficient to pattern tissue deformation and to reduce stresses, including in the ventral, non-remodeling, region. Tissue deformation is thus spatially confined by developmental patterning of the balance between apical and lateral surfaces through membrane recycling, rather than by cytoskeletal forces.

**Highlights:**

- Patterned apical surface expansion in the stretched epidermis confines deformation to the dorsal region of the epidermis
- This pattern arises from membrane recycling, not cytoskeletal reorganization
- Cell flattening under stretch is active process and requires Rab11-dependent recycling
- Dpp/Brinker restricts remodeling dorsally; modeling shows it limits tissue stress

## Introduction

During development, epithelial tissues undergo large deformations, and how they cope with them shapes the final form of the organism (Lecuit and Lenne, 2007; Heisenberg and Bellaïche, 2013). When a tissue is mechanically stretched, it must accommodate by deforming and elongating. Often tissues confine the deformation, and the associated mechanical stresses, to specific regions, so that it occurs where it is needed while the rest of the tissue is preserved. Tissues have been shown to achieve this in several ways. Stretch can trigger local proliferation or oriented cell divisions that progressively relieve stresses by increasing the overall surface (Campinho et al., 2013; Wyatt et al., 2015; Gudipaty et al., 2017), and cell rearrangements can make specific regions of a tissue more fluid, so that they deform while others remain solid (Aigouy et al., 2010; Etournay et al., 2015; Mongera et al., 2018). In these examples, the response of the tissue is mostly attributed to the reorganization of cytoskeletal forces, and the resulting cell shapes, including cell height, are thought to follow passively from the balance of these forces (Hannezo et al., 2014; Sui et al., 2018).

Cells can also respond to stretch by changing their apical surface. To expand its apical surface at constant volume, a cell must become shorter, and its plasma membrane must be redistributed between its apical and lateral sides. Membrane trafficking is known to supply or retrieve membrane during cell shape changes (Sinha et al., 2011; Gauthier et al., 2012; Kosmalska et al., 2015; Figard et al., 2016) and to assist cytoskeletal forces during morphogenesis, for instance during apical constriction (Lee and Harland, 2010; Levayer et al., 2011; Miao et al., 2019). Whether patterned membrane remodeling could provide a way for tissues to confine deformation spatially, and whether developmental signals can control such a pattern, has not been explored.

Dorsal closure of the *Drosophila* embryo offers a clear setting to address these questions (Kiehart et al., 2017; Hayes and Solon, 2017). During closure, the lateral epidermis is stretched dorsally over a hundred of microns by forces generated in the amnioserosa and at the leading edge (Kiehart et al., 2000; Hutson et al., 2003; Solon et al., 2009; Saias et al., 2015; Pasakarnis et al., 2016). Epidermal cells do not divide and undergo minimal intercalation during this process (Kiehart et al., 2017), so the tissue cannot use proliferation or rearrangements to accommodate stretch. The epidermis is also patterned along its dorsoventral axis by Dpp signaling, which is active in its dorsal part and required for epidermal elongation and closure (Glise and Noselli, 1997; Riesgo-Escovar and Hafen, 1997; Fernández et al., 2007; Ducuing et al., 2015). Recent laser-ablation measurements showed that Dpp activity selectively turns the dorsal epidermis from viscoelastic to viscoplastic, allowing it to elongate (Tesson and Vincent, 2025). How this plasticity arises at the cellular level is not known.

Here, we show that, upon stretching, the epidermis confines its deformation to a dorsal region through a patterned increase in apical cell surface. Dorsal cells respond to stretch by expanding their apical surface and becoming shorter. Ventral cells, however, exhibit no change in apical surface area or height. When we increase the stretch on ventral cells, they reorganize their actomyosin like dorsal cells, yet they still do not expand their apical surface or flatten, indicating that the pattern does not originate from cytoskeletal forces. Instead, apical expansion and flattening require active endocytic recycling, through the activity of the small GTPase Rab11. This reveas that changes in cell height under stretch are actively controlled by the cell rather than passively imposed by forces. Using a physical model of the tissue, we show that localized remodeling is sufficient to confine deformation to the remodeling region and to reduce stress in the rest of the tissue. Consistent with this prediction, the Dpp/Brinker pathway restricts remodeling to the dorsal epidermis, and activating Dpp signaling throughout the epidermis spreads deformation to ventral cells. Our results identify patterned membrane recycling as a mechanism by which developmental signaling spatially constrains tissue deformation.

## Results

### Epidermal stretching is spatially patterned during dorsal closure

To follow epidermal deformation during dorsal closure, we imaged embryos expressing DE-Cadherin::Tomato by confocal microscopy from the onset of epidermal stretching (t = 0, 200 min before maximal tissue extension; STAR Methods) until late closure (Figure 1A; STAR Methods and Movie S1). We segmented epidermal cells semi-automatically and assigned each cell to a row according to its position relative to the leading edge of the epidermis (Figure S1B; STAR Methods). For each cell, we measured its length along the DV axis, its width along the anteroposterior (AP) axis and its apical area (Figures 1B, 1C and S1E). We excluded the two rows closest to the leading edge, which have a distinct identity and behavior (Figures S1C and S1D).

**Figure 1.**
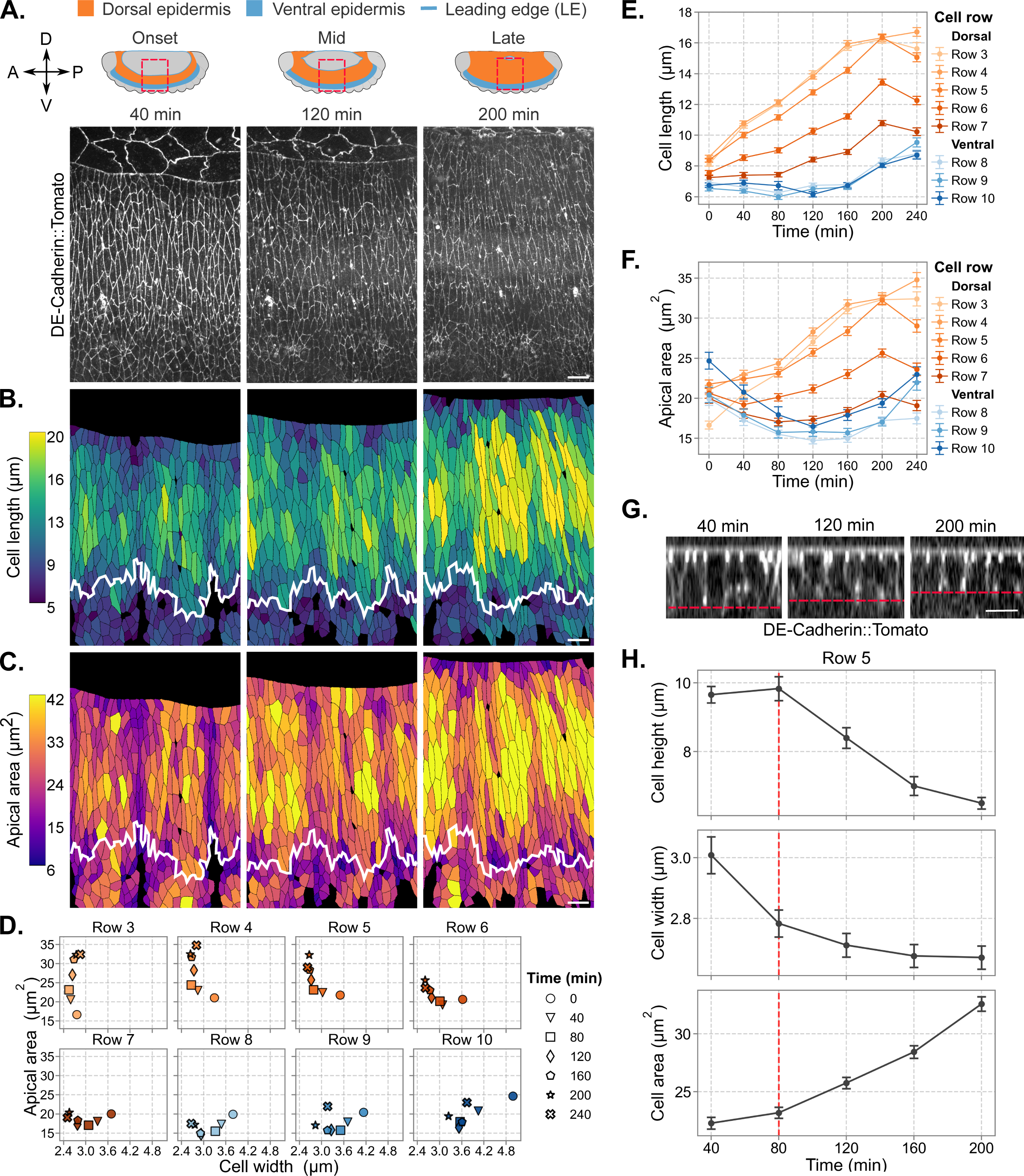
Epidermal stretching is spatially patterned during dorsal closure. (A) Top: schematics of the embryo at the onset, middle and end of dorsal closure (lateral view; anterior to the left, dorsal up), showing the dorsal epidermis (orange), ventral epidermis (blue) and leading edge (LE). The red dashed box indicates the imaged region. Bottom: maximum-intensity projections of DE-Cadherin::Tomato at 40, 120 and 200 min. (B and C) Segmented epidermal cells color-coded by cell length (B) and apical area (C) at the same time points. The white line marks the boundary between dorsal (cell rows 1-7) and ventral epidermis (cell rows 8-10) (D) Apical area versus cell width over time for rows 3–10; symbols indicate time points. (E and F) Cell length (E) and apical area (F) over time for rows 3–10. Orange, dorsal rows; blue, ventral rows. (G) Orthogonal sections of the epidermis (DE-Cadherin::Tomato) in row 5 at 40, 120 and 200 min. The red dashed line indicates the basal surface. (H) Cell height, width and apical area of row 5 cells over time. The red dashed line (80 min) marks the onset of apical area increase and height decrease. Embryos were aligned according to the time of maximal epidermal extension (t = 200 min; see STAR Methods). (E–F) Apical area, cell length and cell width data are presented as mean ± SEM (n=9886, N=6). (H) Height data are presented as mean ± SEM (n=72, N=6). Scale bars: (A–C) 10 µm; (G) 5 µm.

At the tissue scale, we measured the extension of the dorsal domain (P1–P2), the ventral domain (P2–P3) and the whole epidermis (P1–P3) between anatomical landmarks, which we also used to align embryos in time on maximal tissue extension (Figure S1A; STAR Methods). The dorsal domain lengthened by ∼40% between 40 and 180 min, whereas the ventral domain changed little until ∼140–160 min (Figure S1A). At the cell scale, both elongation and apical area increase were spatially heterogeneous (Figures 1B and 1C). Grouping cells by row, we found that cells in dorsal rows 3–5 roughly doubled their length (from ∼8 to ∼16 µm) and their apical area (from ∼17–22 to ∼32 µm²) by 200 min (Figures 1E and 1F). Cells in ventral rows 8–10 elongated much less (from ∼6.5 to ∼8 µm), and their apical area first decreased and then returned close to its initial value. Rows 6 and 7 showed intermediate behavior. Epidermal stretching is therefore spatially patterned, with dorsal cells deforming much more than ventral cells.

### Dorsal cells elongate by expanding their apical surface and reducing their height at constant volume

Cell elongation can result from narrowing at constant apical area, from an increase in apical area at constant width, or from a combination of both (Figure S1H). To distinguish between these modes, we plotted apical area against width over time for each row (Figure 1D). In row 3, cells elongated at nearly constant width (∼2.6–2.8 µm) while their apical area increased. In rows 4–6, cells first narrowed at roughly constant area until they reached a width of ∼2.7 µm, and then expanded their apical area at constant width. In rows 7–10, cells narrowed without sustained increase in apical area (Figures 1D and S1F). Dorsal cells thus elongate in two sequential steps, narrowing followed by apical expansion, whereas ventral cells only narrow.

We next asked whether apical expansion reflects an increase in cell volume or a change in cell shape along the apicobasal axis. We focused on row 5, which switches between the two modes around 80 min (Figures 1D and 1H). Orthogonal sections showed that cell height remained constant (∼9.7 µm) until 80 min and then decreased to ∼6.5 µm at 200 min (Figures 1G and 1H). Over the same period, apical area increased from ∼23 to ∼33 µm², whereas width decreased mainly before 80 min (from ∼3.0 to ∼2.8 µm; Figure 1H). Estimating cell volume as the product of mean height and mean apical area, we found that volume remained approximately constant (∼215 µm³; Figure S1G). Dorsal cells therefore elongate at constant volume, first by narrowing at constant height and then by flattening at nearly constant width, so that apical expansion is compensated by a loss of height.

### Over-stretched ventral cells polarize myosin but neither expand their apical surface nor reduce their height

The patterned deformation could reflect differences in the load experienced by dorsal and ventral cells, or differences in how these cells respond to load. We first compared adhesion and actomyosin organization in the two domains in embryos expressing Sqh::GFP and DE-Cadherin::Tomato (Figure S2A). As previously described (Kaltschmidt et al., 2002; Bulgakova et al., 2013; Gomez et al., 2016), DE-cadherin and myosin II became polarized in opposite manners during elongation, with DE-cadherin enriched at AP-oriented junctions (parallel to the anteroposterior axis of the embryo, i.e., the short cell borders) and myosin II enriched at DV-oriented junctions (parallel to the dorsoventral axis). Both polarizations were stronger in dorsal than in ventral cells (Figures S2B, S2C and Movie S1 and S2). DE-cadherin polarization was weak in ventral cells, whereas myosin remained clearly polarized (∼1.6-fold enrichment). At maximal elongation (200 min), myosin polarization increased sharply in dorsal cells, where myosin formed multicellular cables along DV-oriented junctions (Figures S2A and S2C).

To test whether ventral cells behave differently because they are less stretched, we increased the load on ventral cells by blocking shape changes in the dorsal epidermis. We expressed a constitutively active form of the formin Diaphanous (Dia^CA^) under the control of *pnr*-Gal4, whose expression domain covers rows 1–6 (Figure S2F). Constitutively active Dia increases junctional myosin II and strengthens cell junctions in the embryonic epidermis (Homem and Peifer, 2008; Mulinari et al., 2008), and we reasoned that it would limit the ability of dorsal cells to change shape. Indeed, in *pnr*>Dia^CA^ embryos, the dorsal domain failed to lengthen whereas the ventral domain was over-stretched, and dorsal closure was eventually defective (Figure S2G, and Movie S3). Because perturbed embryos cannot be reliably staged by morphology, we aligned them on the time of maximal extension of the epidermis (P1–P3), defined as t = 200 min, and compared them with controls at this time point and 80 min earlier (t = 120 min; STAR Methods). In these embryos, dorsal cells remained small and not elongated (rows 3–4 at 200 min: length ∼5–6 µm compared with ∼16 µm in controls and apical area ∼12–15 µm², compared with ∼32 µm² in controls; Figures 2A–2E). Ventral cells, in contrast, became more elongated than in controls (rows 8–10 at 200 min: ∼9.7 µm versus ∼8 µm; Figure 2D and Movie S3), and the ventral part of the tissue lengthened more (Figure S2G), indicating that ventral cells were stretched further. These over-stretched ventral cells increased their myosin polarization to levels similar to those of control dorsal cells (Figure S2E and Movie S4), showing that myosin polarization responds to mechanical load regardless of position in the tissue. In contrast, their DE-cadherin polarization remained similar to that of control ventral cells (Figure S2D), even though these cells were more elongated, a shape expected to promote DE-cadherin asymmetry through the alignment of microtubules along the long cell axis (Bulgakova et al., 2013; Gomez et al., 2016).

**Figure 2.**
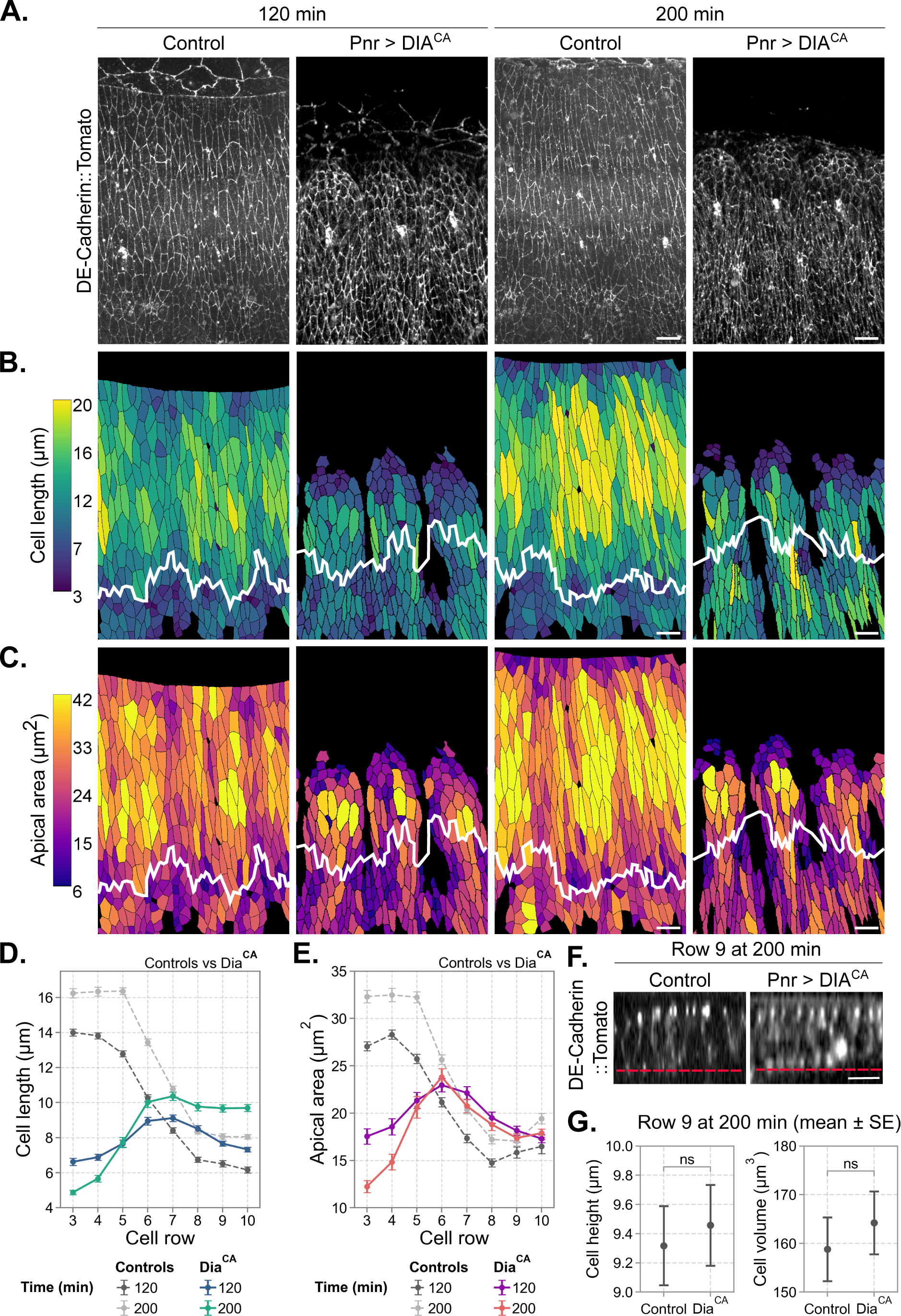
Over-stretched ventral cells neither expand their apical surface nor reduce their height. (A) Maximum-intensity projections of DE-Cadherin::Tomato in control and *pnr*-Gal4 > UAS-Dia^CA^ embryos at 120 and 200 min. (B and C) Segmented cells color-coded by cell length (B) and apical area (C). The white line marks the dorsal–ventral boundary. (D and E) Cell length (D) and apical area (E) per row at 120 and 200 min in control (gray, dashed) and Dia ^CA^ (colored) embryos. (F) Orthogonal sections of row 9 at 200 min in control and Dia^CA^ embryos. The red dashed line indicates the basal surface. (G) Cell height and volume of cells from row 9 at 200 min. Embryos were aligned according to the time of maximal epidermal extension (t = 200 min; see STAR Methods). (D-E) Apical area and cell length data are presented as mean ± SEM (control, n=3399, N=4; Dia^CA^, n= 2403, N=3). (G) Cell height data are presented as mean ± SEM (control, n=11, N=3; Dia^CA^, n=8, N=3). Cell volume was estimated as the product of mean cell height and mean apical area. Statistical comparisons of cell height and volume were performed using a z-test based on group means and standard errors (height, ns; volume, ns). Scale bars: (A–C) 10 µm; (F) 5 µm.

Despite this increased stretch, the apical area of ventral cells in *pnr*>Dia^CA^ embryos remained similar to that of control ventral cells (Figure 2E). They elongated entirely by narrowing, reaching widths of ∼2.4–2.5 µm (Figures S2H-S2I), below the width at which control dorsal cells switch to apical expansion. Their height (∼9.5 µm in row 9) and volume were unchanged compared with controls (Figures 2F-2G) and remained well above the height of control dorsal cells at the same stage (∼6.5 µm in row 5; Figure 1H). Thus, increased stretch is sufficient to elongate ventral cells and to polarize their myosin, but not to trigger apical expansion and height reduction, even when cells become narrower than the width at which dorsal cells switch deformation mode. The ability to expand the apical surface therefore depends on cell identity rather than on the magnitude of stretch.

### Apical expansion and cell flattening require Rab11

Because apical expansion and flattening were restricted to dorsal cells, we asked whether they involve a redistribution of membrane between the lateral and apical domains. To investigate this possibility, we imaged embryos expressing endogenously tagged DE-cadherin-GFP and palmmKate under the control of *pnr*-Gal4, which labels the dorsal domain. Cross-sections of the epidermis showed that, in control embryos, dorsal cells became markedly shorter between early (120 min) and late (200 min) stretching. These cross-sections also showed a reduction of the DE-cadherin signal along their lateral membranes over the same period. In contrast, ventral cells retained prominent lateral E-cadherin signal at 200 min and remained taller (Figure 3A, red arrows). This is consistent with lateral membrane and DE-cadherin being redistributed toward the apical domain as dorsal cells expand their apical surface.

**Figure 3.**
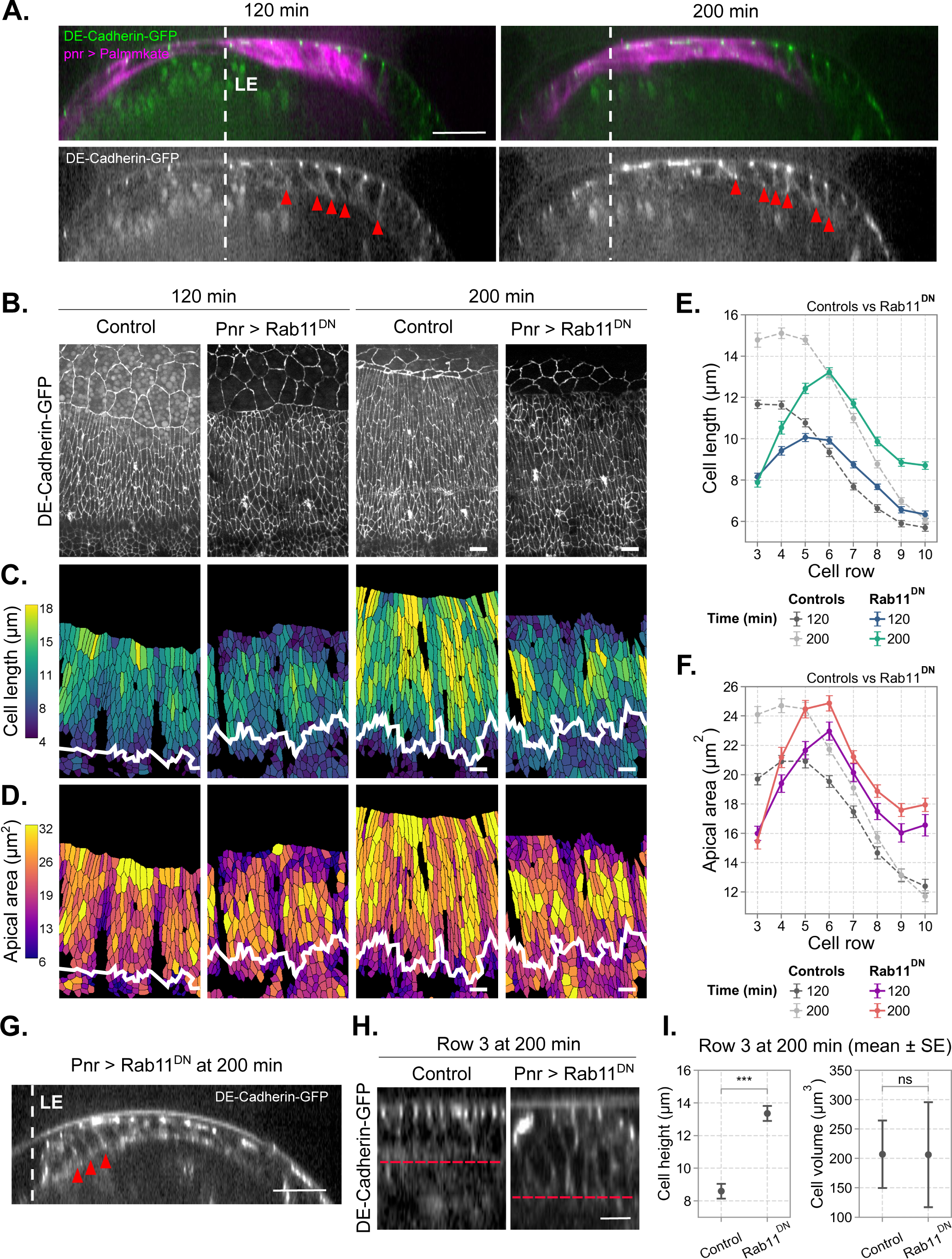
Apical expansion and cell flattening require Rab11. (A) Cross-sections of the epidermis showing cell height and DE-cadherin distribution at early (120 min) and late (200 min) stretching in control embryos expressing *pnr*-Gal4 > UAS-palmmKate. The white dashed line indicates the LE. Note the basolateral junctions in the pannier negative domain (red arrows). (B) Maximum-intensity projections of DE-Cadherin::GFP in control and *pnr*-Gal4 > UAS-Rab11^DN^ embryos at 120 and 200 min. (C and D) Segmented cells color-coded by cell length (C) and apical area (D). The white line marks the dorsal–ventral boundary. (E and F) Cell length (E) and apical area (F) per row at 120 and 200 min in control (gray, dashed) and Rab11^DN^ (colored) embryos. (G.) Cross-section of the epidermis showing cell height and DE-cadherin distribution at late (200 min) stretching in *pnr*-Gal4 > UAS-Rab11^DN^ embryo. The white dashed line indicates the LE. Note the increased cell height and the prominent basolateral junctions in the dorsal domain (red arrows). (H) Orthogonal sections (DE-Cadherin::GFP) of row 3 at 120 and 200 min in control and Rab11^DN^ embryos. The red dashed line indicates the basal surface. (I) Cell height and volume of cells from row 3 at 200 min. Embryos were aligned according to the time of maximal epidermal extension (t = 200 min; see STAR Methods). (E-F) Apical area and cell length data are presented as mean ± SEM (control, n=2791, N=3; Rab11^DN^, n=2867, N=3). (I) Cell height data are presented as mean ± SEM (control, n=11, N=3; Rab11^DN^, n=7, N=3). Cell volume was estimated as the product of mean cell height and mean apical area. Statistical comparisons of cell height and volume were performed using a z-test based on group means and standard errors (height, *** p < 0.001; volume, ns). Scale bars: (A) 20 µm; (B-D) 10 µm; (H) 5 µm.

The endocytic recycling small GTPase Rab11 was shown to be implicated in epidermal elongation during dorsal closure (Mateus et al., 2011). To test whether Rab11 is required for apical expansion, we expressed a dominant-negative form of Rab11 (Rab11^DN^) in the dorsal epidermis using *pnr*-Gal4 (Figure 3B and Movie S5). At 200 min, cells in the most dorsal rows elongated much less than in controls (rows 3–4: ∼8–10.5 µm versus ∼15 µm; Figures 3C and 3E), and their apical area remained low (row 3: ∼15 µm² versus ∼24 µm²; Figures 3D and 3F). They also remained wider than control cells (rows 3–5: ∼2.7 versus ∼2.25 µm; Figures S3A-S3B). The maximum of elongation shifted to rows 5–6 (at the edge of the *pnr* domain, where *pnr* expression decreases; Figure S2F), and ventral rows elongated more than in controls (rows 9–10: ∼9 versus ∼6–7 µm; Figure 3E). At the tissue scale, the dorsal domain no longer lengthened (P1–P2 from ∼55 to ∼48 µm, versus ∼63 to ∼77 µm in controls), whereas the ventral domain lengthened more than in controls (P2–P3 from ∼35 to ∼48 µm, versus ∼31 to ∼40 µm in controls), and the total extension of the epidermis (P1-P3) was reduced (Figure S3C). Ventral rows of Rab11^DN^ embryos already had larger apical areas than controls at 120 min (∼16–17.5 versus ∼12–15 µm²; Figure 3F), which may partly reflect differences in the timing of closure, since embryos were aligned on maximal tissue extension.

Blocking endocytic recycling using Rab11^DN^ also prevented cell flattening. At late stretching, dorsal Rab11^DN^ cells remained tall, and cross-sections indicated that they retained DE-cadherin along their lateral membranes (Figure 3G). At 200 min, cells in row 3 were markedly taller than control cells (∼13.4 versus ∼8.6 µm), whereas their volume was unchanged (∼205 µm³; Figures 3H-3I). Their smaller apical area was therefore compensated by a greater height, consistent with membrane remaining in the lateral domain rather than being redistributed to the apical surface.

Together, these data show that Rab11 is required for apical expansion and flattening. Without it, dorsal cells keep a small apical surface and a large height, and deformation shifts toward ventral rows.

### Localized remodeling is sufficient to confine deformation in a vertex model

To test whether a localized remodeling of cell surfaces is sufficient to confine deformation, we built a two-dimensional vertex model of the apical surface of the epidermis (Farhadifar et al., 2007; Fletcher et al., 2014; Alt et al., 2017; Figures 4A and S4A; STAR Methods). The energy of each cell depends on the deviation of its apical area A and perimeter P from preferred values A_0_ and P_0_, with stiffnesses K_A_ and K_P_, constraining the area and perimeter changes (Figure S4A). The tissue consists of 8 rows of identical cells at rest (With a shape parameter, q_0_ = P_0_/√A_0_ = 4.3, estimated from the shape of epidermal cells at the onset of elongation), corresponding to rows 3–10. The ventral edge is fixed and the dorsal edge is displaced at constant velocity, mimicking the pull of the leading edge and amnioserosa; on the sides, both edges are free to move along the AP axis (Figure 4A). The model describes only the apical surface: the transfer of membrane between the lateral and apical domains is represented by changes in A_0_ and P_0_.

**Figure 4.**
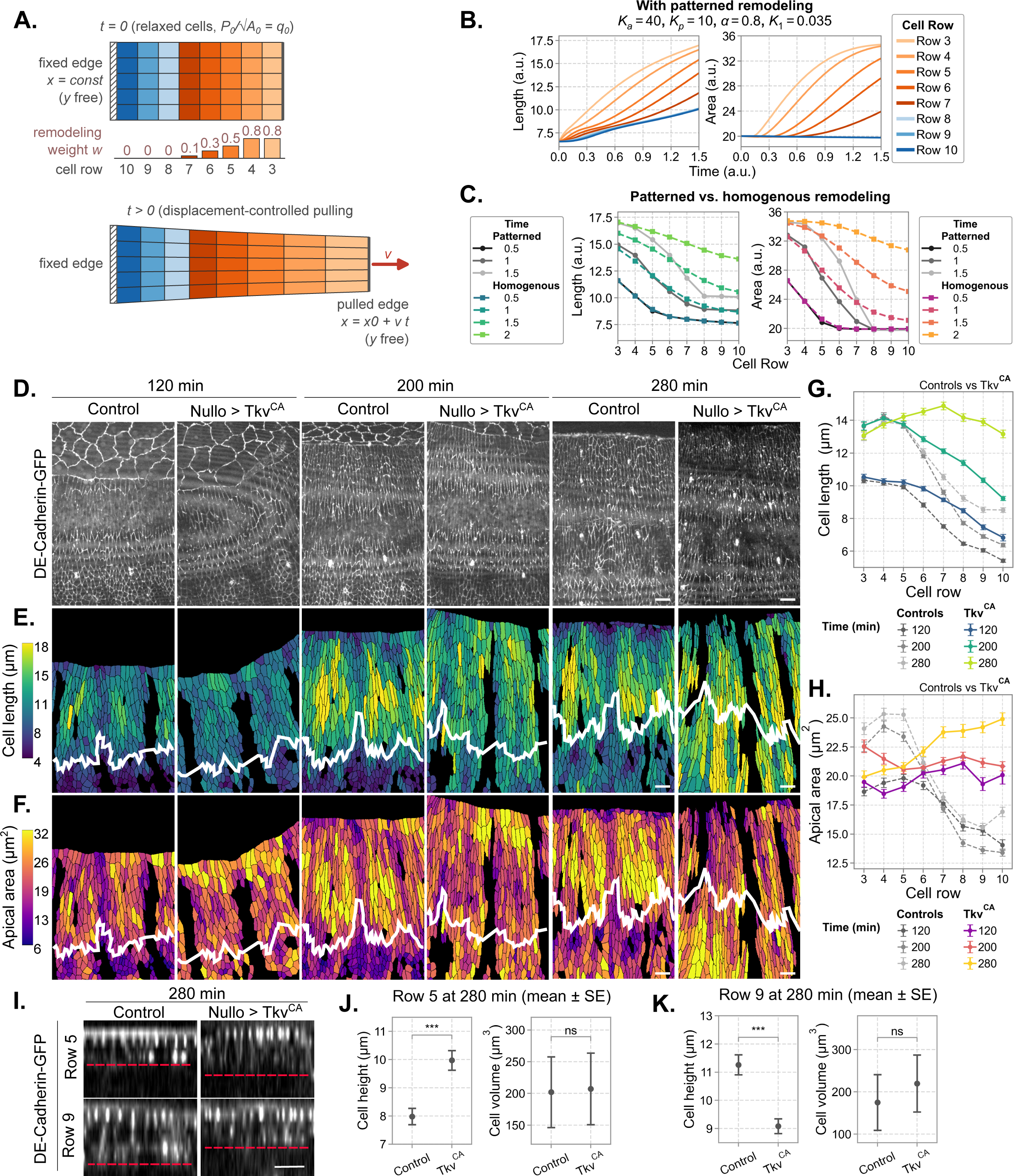
Localized remodeling confines deformation in a vertex model, and uniform Dpp signaling spreads deformation to the ventral epidermis. (A) Representation of the vertex model of the epidermis. The tissue consists of 8 rows of cells (corresponding to rows 3–10), initially at rest with identical preferred area and perimeter (q_0_ = P_0_/√A_0_ = 4.3). The ventral edge is fixed along the DV axis (x) and the dorsal edge is displaced at constant velocity v; both are free along the AP axis (y). Each row was assigned a remodeling weight (w) based on the normalized mean *pnr* expression level per cell row, estimated by measuring the intensity of the palmmKate reporter driven by *pnr*-Gal4 (Figure S2F). (B) Cell length and apical area over time for each row with patterned remodeling (K_A_ = 40, K_P_ = 10, α = 0.8, K_1_ = 0.035). (C) Cell length and apical area per row at successive times with patterned (solid lines) and homogeneous (dashed lines) remodeling. (D) Maximum-intensity projections of DE-Cadherin::GFP in control and *nullo*-Gal4 > UAS-Tkv^CA^ embryos at 120, 200 and 280 min. (E and F) Segmented cells color-coded by cell length (E) and apical area (F). The white line marks the dorsal–ventral boundary. (G and H) Cell length (G) and apical area (H) per row at 120, 200 and 280 min in control (gray, dashed) and Tkv^CA^ (colored) embryos. (I) Orthogonal sections (DE-Cadherin::GFP) of cells from rows 5 and 9 at 280 min in control and Tkv^CA^ embryos. The red dashed line indicates the basal surface. (J and K) Cell height and volume of cells from row 5 (J) at 280 min (K) in control and Tkv^CA^ embryos. Model quantities are in arbitrary units (STAR Methods). Embryos were aligned on maximal extension of the epidermis (t = 200 min; STAR Methods). (G-H) Apical area and cell length data are presented as mean ± SEM (control, n = 4945, N=3; Tkv^CA^, n=4388, N=3). (J-K) Cell height data are presented as mean ± SEM (control row 5, n=9, N=3; Tkv^CA^ row 5, n=11, N=3; control row 9, n=10, N=3; Tkv^CA^ row 9, n=9, N=3). Cell volume was estimated as the product of mean cell height and mean apical area. Statistical comparisons of cell height and volume were performed using a z-test based on group means and standard errors (height row 5, *** p < 0.001; volume row 5, ns; height row 9, *** p < 0.001; volume row 9, ns). Scale bars: (D-F) 10 µm; (I) 5 µm.

Without remodeling, cells elongated by narrowing, and their apical area remained constant or decreased (Figure S4C). Depending on K_P_, elongation was either nearly uniform across rows (K_P_ = 10) or graded from dorsal to ventral (K_P_ = 1), because deformation propagates more slowly into a softer tissue. Interestingly, without remodeling we don’t observe apical area increase independently of the parameters (Figure S4C). Passive mechanics can therefore reproduce a transient gradient of cell length, which disappears as the tissue equilibrates, but not the dorsal increase in apical area observed in embryos.

We then allowed cells to remodel their preferred shape when stretched. When the perimeter of a cell exceeds its preferred value by more than a threshold ε, its preferred area grows at a rate proportional to the excess strain, up to a maximal value A_0f_, and its preferred perimeter grows in proportion, with a ratio α between perimeter and area increments (Figure S4B; STAR Methods). This represents the irreversible addition of membrane to the apical surface. We weighted the remodeling rate of each row by the average *pnr*-expression in cells measured in that row (w = 0.8 for rows 3–4, 0.5 for row 5, 0.3 for row 6, 0.1 for row 7 and 0 for rows 8–10; Figures 4A and S2F). All parameters except the remodeling rate were constrained by experimental observations (STAR Methods): the initial and maximal preferred areas match the initial and final apical areas of dorsal cells, the partition of added membrane between area and perimeter (α) sets the final width of remodeling cells, and the remodeling threshold ε reproduces the delay between the increase in perimeter and the increase in area observed in dorsal cells (Figure 1H). With this patterned remodeling, dorsal rows elongated and increased their apical area, whereas ventral rows elongated less and kept a constant area (Figure 4B and Movie S6), recapitulating the behavior of the embryonic epidermis (Figures 1E and 1F). The model reproduces the pattern qualitatively but not every feature of the data: ventral cells elongate somewhat more than in embryos, and their area does not show the transient decrease observed *in vivo*.

We next asked how the spatial pattern of remodeling affects the tissue. When all rows were given the same remodeling weight, deformation and apical area increase were no longer confined to the dorsal rows but spread into the ventral rows (Figure 4C and Movie S7). A gradient remained, because stretch is transmitted from the pulled edge, but ventral cells elongated more and increased their apical area. Localized remodeling therefore confines deformation to the remodeling region. To assess how remodeling affects stress in the tissue, we computed the perimeter tension of each cell, 2K_P_(P − P_0_), with and without remodeling, using identical mechanical parameters (Figure S4D). In both cases, tension was highest in the row closest to the pulled edge and decreased toward the fixed edge. Without remodeling, tension increased steadily in all rows. With patterned remodeling, it built up more slowly and, at the end of the simulation, was ∼40–50% lower in all rows (∼280 versus ∼465 a.u. in row 3), with the largest reduction in the non-remodeling ventral rows (∼190 versus ∼365 a.u. in row 10). Localized remodeling thus confines deformation to the remodeling region while reducing the stress transmitted to the rest of the tissue. Remodeling raised the target shape index q_0_ = P_0_/√A_0_ of remodeling rows from 4.3 to up to ∼5.3, whereas it remained at 4.3 in non-remodeling rows (Figure S4D). The shape index indicates the fluidity/plasticity of the tissue, in case of rectangular cells a shape index of 4 indicates a solid tissue and increase in this index indatcates a potential fluidization of the tissue (Bi et al., 2015). Here, this growth of preferred perimeter relative to preferred area lowers the energetic cost of elongation, so remodeling rows deform plastically, whereas non-remodeling rows deform only elastically and resist further stretching. Cell rearrangements were not included in the model, as they are rare in the epidermis during dorsal closure; the plastic deformation of remodeling rows therefore does not involve tissue flow.

### Dpp/Brinker signaling restricts surface remodeling to the dorsal epidermis

The model predicts that extending remodeling to the whole tissue should spread deformation and apical area increase to ventral cells. In the embryo, the remodeling domain coincides with the dorsal domain of *pnr* expression, where Dpp signaling is active (Glise and Noselli, 1997; Riesgo-Escovar and Hafen, 1997; Figures S2F, S4F). Mutants affecting Dpp signaling show defects in epidermal elongation and dorsal closure (Fernández et al., 2007; Ducuing et al., 2015), and early Dpp activity is required for the plastic deformation of the dorsal epidermis during closure (Tesson and Vincent, 2025). To test whether Dpp signaling defines the remodeling domain, we expressed a constitutively active form of the Dpp receptor Thickveins (Tkv^CA^) throughout the epidermis using *nullo*-Gal4. The Dpp reporter Dad-RFP (Tsuneizumi et al., 1997), enriched in dorsal cells in controls, was expressed throughout the epidermis in these embryos (Figure S4G).

As predicted by the model, Tkv^CA^ homogeneous expression in the epidermis spreads deformation across the epidermis (Figures 4D–4H and Movie S8). In Tkv^CA^ expressing embryo, the overall tissue distances (P1–P3) are more extended, supporting the confinement argument (Figure S4E). At 200 min, ventral cells (rows 8–10) were more elongated than in controls (∼9–12 µm versus ∼6.5–7.5 µm) and had a larger apical area (∼21 µm² versus ∼14– 16 µm²); by 280 min, their apical area (∼24–25 µm²) exceeded that of dorsal cells (Figures 4G-4H). Dorsal cells, in contrast, increased their apical area less than in controls (∼20–21 versus ∼24–25 µm² at 280 min; Figure 4H), consistent with tissue extension being shared over more cells. Cells in all rows were also narrower than in controls (Figures S4H and S4I). Consistent with a more uniformly distributed deformation across the epidermis, at 280 min, dorsal and ventral cells exhibited more homogeneous heights in Tkv^CA^ embryos (∼9.94 versus ∼9.05 µm), whereas ventral cells were taller than dorsal cells in control embryos (∼7.85 versus ∼11.12 µm; Figures 4I-4K). In both cases cell volume remained constant (∼200 µm³; Figures 4J-4K). Absolute values in controls differ from those in Figure 1, likely because of differences in genotype. The patterned behavior is, however, conserved, and each genotype is therefore compared with matched controls.

Conversely, we asked whether repressing Dpp target genes in the dorsal epidermis is sufficient to suppress remodeling. The Dpp is spatially constrained by Brinker (Brk), a transcriptional repressor of Dpp targets, expressed where Dpp signaling is low (Campbell and Tomlinson, 1999; Jaźwińska et al., 1999; Minami et al., 1999; Hamaratoglu et al., 2014; Figure S5L). We forced the expression of Brk in the dorsal epidermis using *pnr*-Gal4 to identify whether Brk impact the ability of cells to elongate. The Dpp pathway reporter Dad-RFP showed a significant decrease in fluorescence intensity (Figure S5K), consistent with Brk-mediated repression of Dpp target genes. In Brk overexpressing embryos dorsal cells failed to elongate (rows 3–5: ∼10.5–11 µm at both 120 and 200 min, whereas control cells elongated from ∼10.5 to ∼14 µm), and their apical area decreased instead of increasing as they narrowed (from ∼24 to ∼19–20 µm², whereas control cells increased from ∼19 to ∼23–24 µm²; Figures S5A-B–G). At the tissue scale, the dorsal domain shortened instead of lengthening (P1–P2 from ∼50 to ∼39 µm, versus ∼63 to ∼77 µm in controls), and the total extension of the epidermis was reduced (Figure S5J and Movie S9). Brk embryos already differed from controls at 120 min, including in ventral rows, which may partly reflect differences in the timing of closure. At 200 min, dorsal cells were taller in Brk overexpressing embryos than in controls (∼10.44 versus ∼7.8 µm; Figures S5H-S5I), while cell volume remained constant (∼200 µm³; Figure S5I).

Together, these results show that Dpp/Brinker signaling defines the domain in which cells remodel their surface under stretch. Activating Dpp signaling throughout the epidermis spreads remodeling and deformation to ventral cells, as predicted by the model, whereas Brinker expression in dorsal cells suppresses them (Figure S5L).

## Discussion

Developing tissues must accommodate to large stretch by elongation and deforming. However, confining the deformation and stresses to the regions close to the stretch is essential to protect tissue integrity. Our study identifies a new way of achieving this. During dorsal closure, the epidermis confines its deformation to a dorsal region through a patterned increase in apical cell surface, driven by membrane recycling and restricted to this region by the Dpp/Brinker pathway.

### Patterned remodeling confines deformation and limits stress

Stretched epithelia have been shown to relieve stress through local proliferation and oriented divisions (Campinho et al., 2013; Wyatt et al., 2015; Gudipaty et al., 2017) or through cell rearrangements that fluidize specific regions (Bi et al., 2015; Mongera et al., 2018). The dorsal closure epidermis uses neither of these methods. Our data and model point to a third strategy: cells in a defined region remodel their surface, which allows them to change their resting shape and deform permanently, without dividing or exchanging neighbors. In the model, restricting this remodeling to the dorsal region is sufficient to concentrate deformation there and to reduce the stress transmitted to the rest of the tissue, including the non-remodeling rows. The embryo behaves as the model predicts: when Dpp signaling is activated throughout the epidermis, deformation spreads into the ventral epidermis and dorsal cells deform less, and when Rab11^DN^ or Brinker expression in dorsal cells blocks remodeling in dorsal cells, the dorsal domain no longer lengthens and deformation shifts toward the ventral epidermis. The dorsal epidermis therefore acts as a deformable zone that absorbs the stresses by a localized stretch, protecting the ventral epidermis.

### A patterned response to stretch that does not come from cytoskeletal forces

A natural explanation for the patterned deformation of the epidermis would be that dorsal and ventral cells experience different forces, or organize their cytoskeleton differently. Our data argue against this. When ventral cells are over-stretched, they polarize their actomyosin to the same extent as dorsal cells, showing that the cytoskeleton responds to stretch everywhere in the tissue. Yet these cells still do not expand their apical surface. The cytoskeletal response is therefore not what distinguishes dorsal from ventral cells. What distinguishes them is their ability to remodel their surface, which depends on Rab11 and is lost when dorsal cells express Brinker. This places membrane recycling, rather than force generation, at the origin of the pattern. Membrane trafficking has so far mostly been seen as supporting force-driven shape changes, for instance during apical constriction (Lee and Harland, 2010; Levayer et al., 2011; Miao et al., 2019) or by buffering membrane tension (Gauthier et al., 2012; Kosmalska et al., 2015; Figard et al., 2016). Our results show that it can instead determine where a tissue deforms.

### Cell height is actively controlled under stretch

A purely mechanical view predicts that a stretched cell flattens passively, its height adjusting to the tensions of its apical and lateral surfaces (Hannezo et al., 2014; Sui et al., 2018). Our observations do not fit this view. Over-stretched ventral cells become longer and narrower but keep their height, and dorsal cells flatten only when Rab11 is functional. Flattening therefore requires the cell to actively move membrane from its lateral to its apical side. Forces may set the shape of the apical surface, but how a cell distributes its membrane between the apical and lateral sides, and thus how tall it is, is controlled by recycling. This suggests that the three-dimensional shape of epithelial cells, and of the tissues they form, can be tuned by regulating membrane traffic, independently of changes in force.

### Developmental signaling patterns the mechanical response of a tissue

Dpp/Brinker signaling is best known for patterning tissues and controlling their growth, with Brinker repressing Dpp signaling and constraining Dpp expression domain (Campbell and Tomlinson, 1999; Jaźwińska et al., 1999; Minami et al., 1999; Schwank et al., 2008; Restrepo et al., 2014). Tesson and Vincent (2025) recently showed that Dpp activity impacts dorsal epidermis mechanics, making it viscoplastic, so that it deforms permanently under stretch. Our results identify a cellular mechanism for this plasticity: cells with high Dpp signaling remodel their apical and lateral surfaces under stretch, whereas Brinker-expressing cells do not, and our model shows that such localized remodeling is sufficient to produce plastic, spatially confined deformation. Dpp signaling was already known to influence epithelial cell shape, promoting the columnar shape of wing disc cells (Widmann and Dahmann, 2009) and the maintenance of cells within the wing disc epithelium (Gibson and Perrimon, 2005; Shen and Dahmann, 2005). We identify membrane recycling as a way in which Dpp signaling shapes cells. Trafficking is also known to shape Dpp signaling itself (Entchev et al., 2000), suggesting that the two processes may be closely interlinked. Because BMP signaling also regulates adhesion and cell shape during vertebrate morphogenesis, for example during gastrulation (von der Hardt et al., 2007) and neural tube closure (Eom et al., 2011), patterned control of membrane recycling by BMP signaling may be a general way for developing tissues to confine deformation.

## Materials and Methods

### Fly strains and husbandry

*Drosophila melanogaster* stocks were raised and crossed at 22°C on standard cornmeal–agar medium (composition in the Key resources table). All strains used are listed in the Key resources table.

Embryos expressing DE-cadherin::Tomato and Sqh::GFP were used to characterize the process in Figure 1 and as controls in Figure 2. For Figure 2, Dia^CA^ was expressed under the control of *pannier*-Gal4 (*pnr*-Gal4) in embryos expressing DE-cadherin::Tomato and Sqh::GFP.

In Figures 3 and S5, embryos expressing endogenously tagged DE-cadherin-GFP and *pnr*-Gal4 were crossed with flies carrying UAS-Rab11^DN^ (Figure 3) and UAS-Brk (Figure S5). For the controls, these flies were crossed with UAS-palmmKate. The palmmKate reporter was used to identify embryos expressing *pnr*-Gal4, which were heterozygous for the Gal4 driver. The effectiveness of Brk overexpression in the dorsal domain was assessed crossing UAS-Brk with *pnr*-Gal4 flies carrying an endogenously tagged Dad-RFP, reporter of Dpp pathway activity.

For Figure 4, UAS-Tkv^CA^ was expressed under the control of *nullo*-Gal4 in embryos expressing endogenously tagged DE-cadherin-GFP and Dad-RFP. As UAS-Tkv^CA^ expression was heterozygous, Dad-RFP fluorescence was used to identify embryos in which Tkv^CA^ overexpression was effective. Embryos exhibiting ectopic Dad-RFP signal in ventral cells, in addition to the dorsal signal, were classified as Tkv^CA^-overexpressing embryos, whereas embryos displaying the wild-type dorsal-restricted Dad-RFP pattern were used as controls.

### Embryo collection and mounting

Stage 12-15 embryos were collected at 22°C on molasses plates (composition in the Key resources table) and aged overnight at 18°C. They were dechorionated in 5% sodium hypochlorite for 2 min, rinsed in tap water for 1 min, mounted dorso-laterally (lateral epidermis facing the coverslip) on a coverslip coated with heptane glue, and covered with halocarbon oil to prevent desiccation.

### Live imaging

Time-lapse z-stacks were acquired on an Olympus IXplore SpinSR10 spinning-disk confocal microscope with a 60×/1.42 UPLXAPO oil or a 100×/1.50 UPLAPO oil objective, using 2×2 binning, resulting in pixel sizes of 0.216 µm and 0.13 µm, respectively. The z-step was 1 µm, except for the Tkv^CA^, and corresponding control experiments, for which it was 2 µm. Stacks were acquired every 10 min (Rab11^DN^, Dia^CA^, Tkv^CA^ and Brk experiments) or 20 min (controls in Figures 1 and 2). GFP was excited with the 488 nm laser line and Tomato and RFP/mKate with the 561 nm line. All experiments were performed at 22°C, except the Rab11^DN^ experiments, which were imaged at 25°C.

### Image processing

Images were processed in Fiji (Schindelin et al., 2012) and analyzed with custom Python code. Unless otherwise stated, analyses were performed on maximum-intensity projections of z-stacks after background subtraction and bleach correction in Fiji (exponential correction).

### Cell segmentation and shape measurements

The apical surface of epidermal cells was segmented semi-automatically on DE-cadherin images using a fine-tuned, pre-trained Cellpose model (Stringer et al., 2021; Pachitariu and Stringer, 2022), followed by manual correction. Each cell was manually assigned to a row: cells in contact with the leading edge were assigned to row 1, and row numbers increased ventrally (Figure S1B).

For each cell, using a custom python code, we measured the apical area and fitted an ellipse to its outline; cell length and width were defined as the lengths of the major and minor axes of this ellipse, and the aspect ratio as length divided by width. Major axis is aligned with the DV axis in most cells, whereas the minor axis in aligned with the AP axis.

Because the embryo is curved along its DV axis, projecting it onto a two-dimensional image underestimates distances along this axis, increasingly so with distance from the region in contact with the coverslip. We corrected for this by assuming that the projection error arises from the curvature along the DV axis. For each cell, we computed a corrected pixel size in the DV direction that depends on the cell’s position along the DV axis, accounting for the real shape of the embryo along DV axis, resulting in rectangular pixels, and used it to correct area, length and width to account of the curvature.

### Junction polarization

Because the leading edge is curved, the DV axis differs between segments. For each segment, we defined the local DV axis as a straight line from the leading edge to the ventral edge of the epidermis. For each cell edge, we measured its angle to the DV axis of its segment and classified it as parallel (0–20°) or perpendicular (30–90) to the DV axis. Mean DE-cadherin and myosin II intensities were measured for each edge using a dilated edge mask.

For each cell, the DE-cadherin polarization ratio was defined as the mean DE-cadherin intensity at perpendicular edges divided by that at parallel edges, so that a ratio above 1 indicates enrichment at edges perpendicular to the DV axis. Because myosin II is expected to show the opposite enrichment, its polarization ratio was defined as the mean intensity at parallel edges divided by that at perpendicular edges, so that a ratio above 1 indicates enrichment at edges parallel to the DV axis.

### Cell height and volume

Orthogonal sections of selected cell rows were generated from DE-cadherin images using the *Reslice* function in Fiji. Cell height was measured manually by drawing apical-to-basal lines along the cell junctions visible in the orthogonal sections. For each embryo, 2–4 segments were analyzed, with 3–5 embryos analyzed per condition. The sample size (n) for the height analysis corresponds to the mean height value calculated per segment.

Mean cell volume was estimated as the product of mean apical area and mean cell height, assuming a prismatic cell geometry. The SEM of the estimated mean volume was calculated using first-order error propagation for the product of two variables, assuming that apical area and cell height were independent.

### Tissue-scale measurements and time alignment

Tissue-scale deformation was quantified as the distances between three landmarks along the DV axis: P1, located at the leading edge; P2, corresponding to the third sensillum at the boundary between the dorsal and ventral domains; and P3, corresponding to the fourth sensillum in the ventral epidermis. The P1–P2 distance represents the dorsal domain, P2–P3 the ventral domain, and P1–P3 the maximal epidermal extension (Figure S1A). Distances were measured in Fiji, every 10 min, along a line following the DV axis of each segment. For each embryo, 2–4 segments were analyzed, with 3–5 embryos analyzed per condition.

To compare embryos at equivalent stages, embryos were aligned in time on the maximal extension of the epidermis (P1–P3 distance; Figure S1A). This distance increases during closure until it reaches a maximum, which was set to t = 200 min; t = 0 therefore corresponds to 200 min before maximal tissue extension. In control embryos this maximal extension starts reducing shortly after this maximum (t = 240 min). Because perturbed embryos could not be reliably staged by morphology, the same criterion was used for all genotypes, and perturbed embryos were compared with controls at t = 200 min and 80 min earlier (t = 120 min). In the case of Dia^CA^, Rab11^DN^ and Brk, in which a clear maximum was not observed, the reference time point was defined as the point at which the P1–P3 distance curve reached a plateau.

### Statistical tests

For each condition in cell shape plots and in polarization ration plots, N refers to the number of embryos and n to the total number of cells analyzed; these values are indicated in the figure legends. Data are presented as mean ± SEM.

For comparisons of polarization ratios, the statistical significance was assessed using two-way ANOVA followed by Benjamini-Hochberg corrected Welch’s t-tests.

For comparisons of cell height and cell volume between control and mutant embryos, a z-test based on group means and standard errors was used. N refers to the number of embryos and n to the segments analyzed. These values are indicated in the figure legends. Data are presented as mean ± SEM.

### Vertex model

#### Geometry and energy

We modeled the apical surface of the epidermis as a two-dimensional vertex model (Farhadifar et al., 2007; Fletcher et al., 2014; Alt et al., 2017). The tissue consisted of 8 rows of cells along the DV axis, corresponding to epidermal rows 3–10, and 5 cells along the AP axis (Figure 4A). Each cell is a quadrilateral defined by its four vertices, and adjacent cells share vertices. The mechanical energy of the tissue is

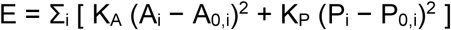

where A_i_ and P_i_ are the area and perimeter of cell i, A_0,i_ and P_0,i_ its preferred area and perimeter, and K_A_ and K_P_ the area and perimeter stiffnesses, identical for all cells (Figure S4A). Areas were computed with the shoelace formula.

#### Dynamics

Vertex positions r_k_ evolve according to overdamped dynamics,

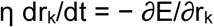

with a friction coefficient η identical for all vertices. Equations were integrated numerically using explicit Euler with a time step of Δt=2.10^-6^, chosen small enough that results did not change when it was halved.

#### Initial state and boundary conditions

All cells initially had identical preferred area and perimeter, with q_0_ = P_0_/√A_0_ = 4.3, estimated from the shape of epidermal cells at the onset of elongation, and A_0_ = 20 (arbitrary units). The initial configuration was obtained by minimizing the energy without external load, so that the simulations start from mechanical equilibrium. The vertices of the ventral edge (row 10) were then held at a fixed position along the DV axis (x), and those of the dorsal edge (row 3) were displaced along x at constant velocity v, x(t) = x_0_ + vt, mimicking the pull exerted by the leading edge and the amnioserosa. Both edges were free to move along the AP axis (y), and the two AP boundaries of the tissue were free (Figure 4A).

#### Remodeling

Remodeling represents the irreversible addition of membrane to the apical surface of stretched cells. When the perimeter of a cell exceeds its preferred value by more than a threshold ε, the cell experiences a strain

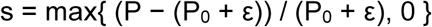

and its preferred area and perimeter increase as

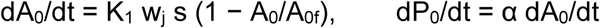

where K_1_ sets the remodeling rate, A_0f_ is the maximal preferred area (common to all cells), α is the ratio between perimeter and area increments of the added membrane, and w_j_ is the remodeling weight of row j (Figure S4B). Because s ≥ 0 and A_0_ ≤ A_0f_, remodeling is irreversible and saturating. For patterned remodeling, w_j_ was set to the fraction of *pnr*-expressing cells measured in the corresponding row (Figure S2F): w = 0.8 for rows 3 and 4, 0.5 for row 5, 0.3 for row 6, 0.1 for row 7 and 0 for rows 8–10 (Figure 4A). For homogeneous remodeling, all rows were given the same weight, w = 0.8. Simulations without remodeling used w = 0 for all rows.

#### Parameters

Model parameters were constrained by experimental observations wherever possible. Lengths are expressed in units such that the initial preferred area (A_0_ = 20) matches the initial apical area of epidermal cells (∼17–25 µm²; Figure 1F), so that one length unit corresponds to approximately 1 µm. During early elongation, dorsal cells increase their perimeter at nearly constant apical area (row 5 between 40 and 80 min; Figure 1H), indicating that area changes are more costly than perimeter changes; we therefore set K_A_ > K_P_. Their values (K_A_ = 40, K_P_ = 10) were chosen to reproduce the delay in elongation observed between dorsal and ventral rows (Figure 1E). The pulling velocity v was set to reproduce the total deformation of rows 3–10 (Figure 1E), so that the total extension of the tissue over the simulation matched the measured extension of these rows. The maximal preferred area A_0f_ = 35 was set to the apical area reached by row 3 at the end of elongation, where its increase levels off (Figure 1F). The parameter α, which sets how added membrane is partitioned between preferred area and perimeter, determines the width that cells reach after remodeling; we set α = 0.8 so that the final width of remodeling cells matched that of dorsal rows (∼2.6– 2.8 µm; Figure S1F). α cannot be read directly from the measured increments of area and perimeter, because the measured perimeter of stretched cells deviates from their preferred perimeter. The threshold ε = 1.8 was chosen to reproduce the temporal shift observed in dorsal cells between the increase in perimeter and the later increase in apical area (Figure 1H). The remodeling rate K_1_ = 0.035 is the only parameter not constrained by experiment. It was previously selected by fitting the experimental dA–dP relationship and the increase in area. Because time is in arbitrary units, K_1_ sets how fast remodeling proceeds relative to tissue stretching; varying it changed how quickly remodeling cells reached A_0f_, but not their final area or the confinement of area increase to the remodeling rows. Simulations without remodeling (Figure S4C) used (K_A_, K_P_) = (10, 10), (40, 10) and (40, 1).

For each cell, we computed its area, perimeter, length and width, and averaged them per row. For the stress readout: per-cell perimeter tension Λ_i_ = 2K_P_(P_i_ − P_0,i_), averaged per row; and the target shape index p_0_ = P_0_/√A_0_ per row. The model was implemented in Python using NumPy and SciPy (energy minimization).

## Key resources table

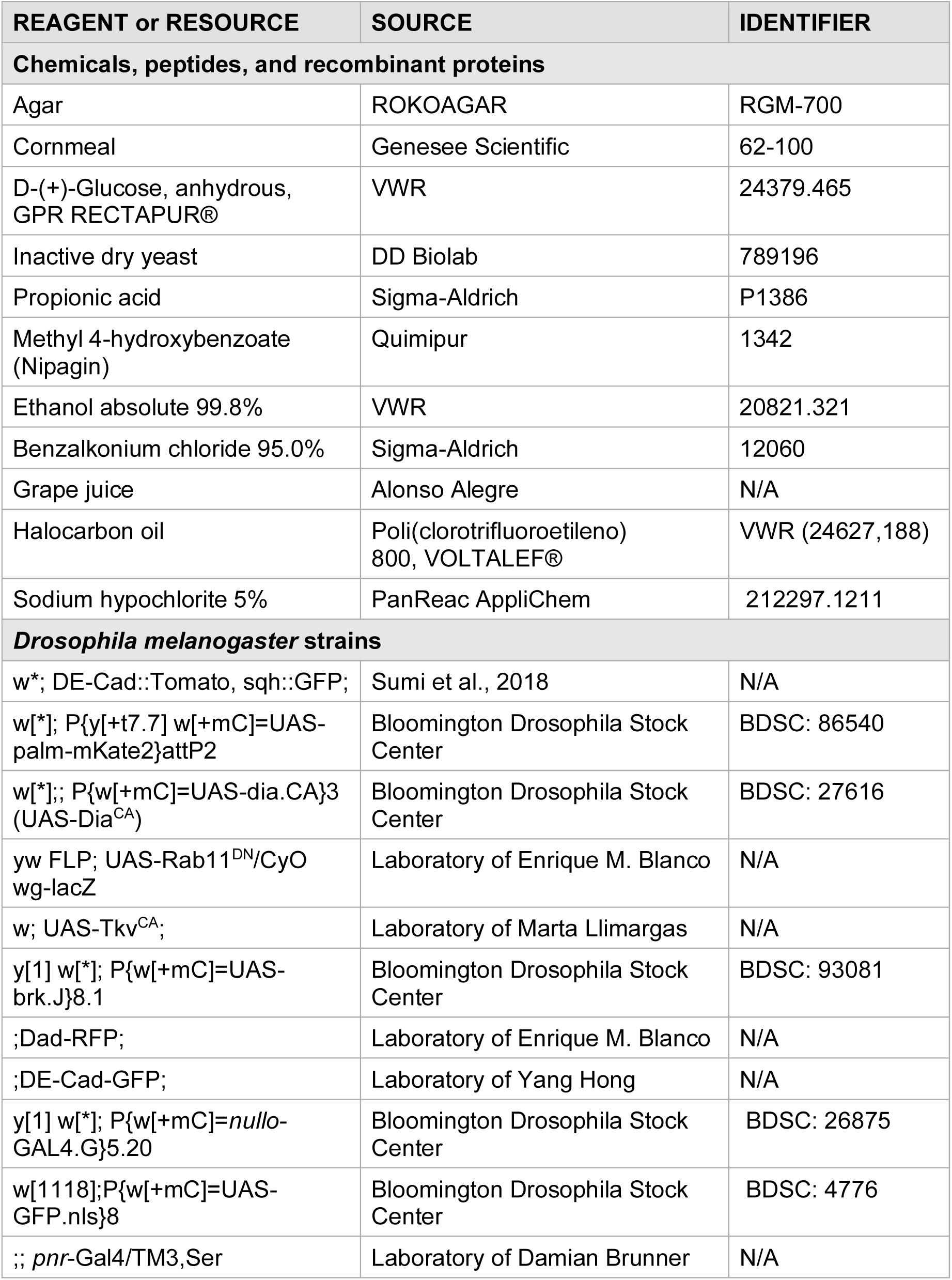

## Acknowledgements

We thank Marta Llimargas and Enrique Martin Blanco for providing flie lines. We thank the BRALM microscopy core facility. The research leading to these results has received funding from the Spanish Ministry of Science and Innovation, Plan National, PID2019-109117GB-100, PID2022-142779NB-100 and from the Basque government, Proyectos de Investigacion Basica/Aplicada, PIBA_2023_1_0033. This work was supported in part by the Fundación Biofisika Bizkaia.

