## Supplemental figures and legends for "Patterned membrane recycling balances apical and lateral cell surfaces to confine tissue deformation under stretch"

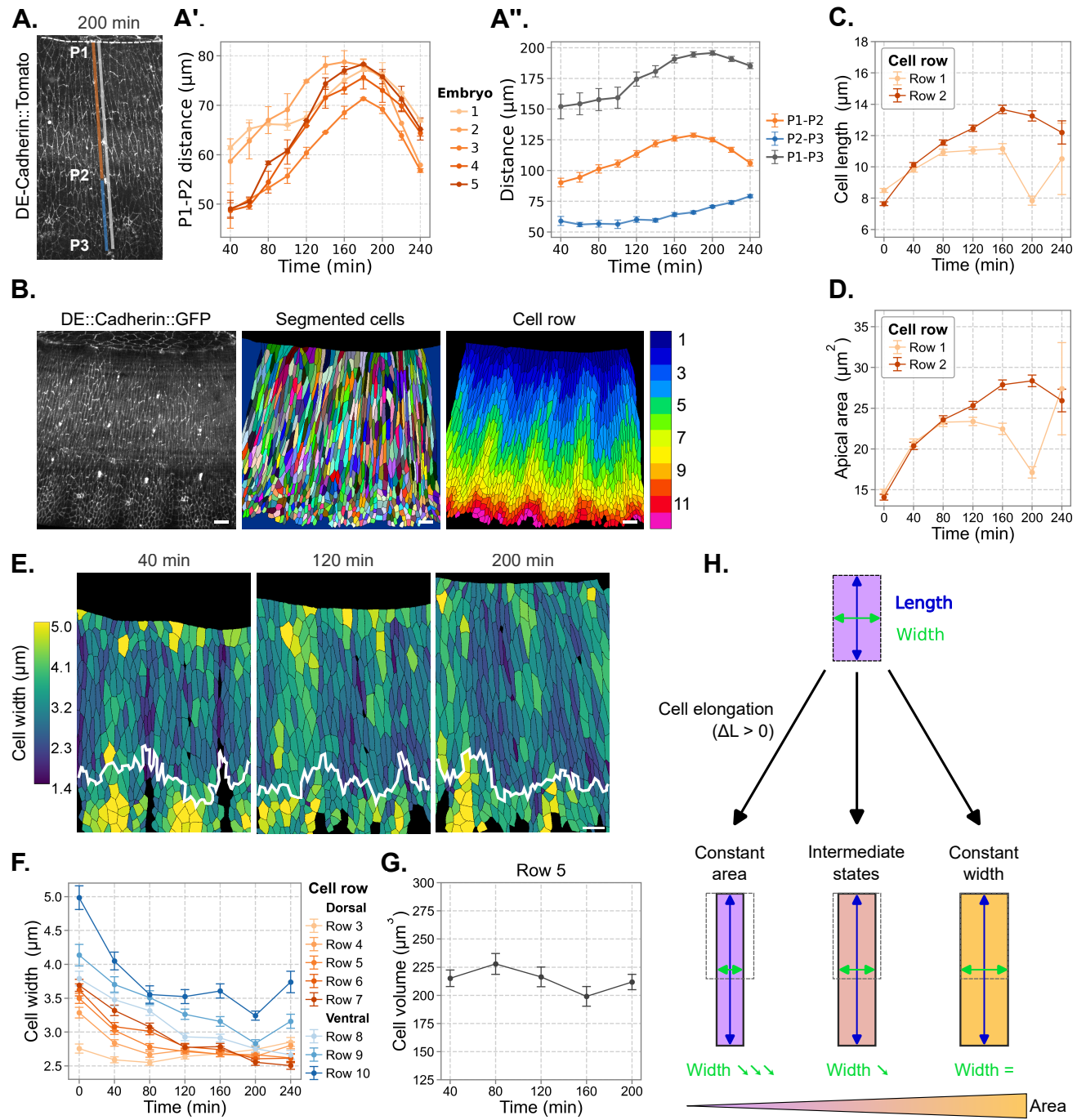

**Figure S1. Tissue-scale and cell-scale deformation of the epidermis in control embryos, related to Figure 1**

(A) DE-Cadherin::Tomato at 200 min showing landmarks P1 (leading edge), P2 (third sensillum) and P3 (fourth sensillum). Orange, blue and gray lines represent respectively dorsal domain (P1-P2 distance), ventral domain (P2-P3 distance) and total epidermis extension (P1-P3 distance). (A') Alignment of individual embryos based on dorsal domain extension (P1-P2) over time ($n = 5$). (A'') Mean length of the dorsal domain (P1-P2), ventral domain (P2-P3), and total epidermis extension (P1-P3) over time ($N = 5$). Note that maximum epidermal extension (P1-P3) is reached at 200 min. (B) Image processing pipeline: original DE-Cadherin::GFP image, segmented cells, and cells color-coded by row. (C and D) Cell length (C) and apical area (D) over time for rows 1 and 2. (E) Segmented cells color-coded by cell width at 40, 120 and 200 min. The white line marks the dorsal-ventral boundary. (F) Cell width over time for rows 3-10. (G) Volume of row 5 cells over time. (H) Possible modes of cell elongation: at constant apical area (by narrowing), at constant width (by apical expansion), or intermediate states combining both.

(C-D) Apical area and cell length data are presented as mean ± SEM ($n = 2358$, $N=6$). (F) Cell width data are presented as mean ± SEM ($n=9896$, $N=6$). (G) Cell volume was estimated as the product of mean cell height and mean apical area. Scale bars: (B, E) 10 $\mu\text{m}$.

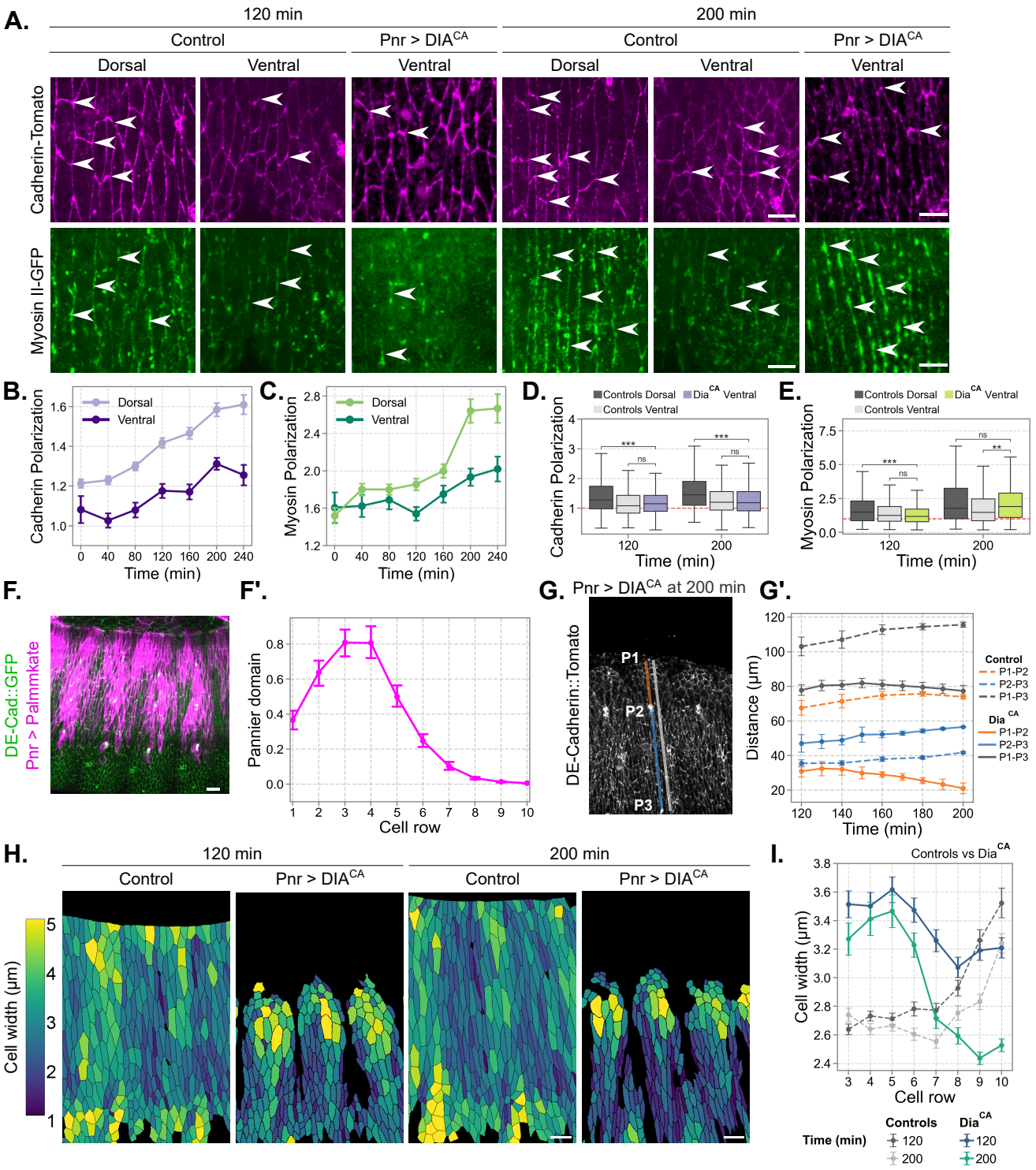

**Figure S2. DE-Cadherin and myosin II polarization, pannier domain and tissue deformation in *Dia<sup>CA</sup>* embryos, related to Figure 2**

(A) DE-Cadherin::Tomato (magenta) and myosin II (*Sqh*::GFP, green) in dorsal and ventral regions of control embryos and in the ventral region of *Dia<sup>CA</sup>* embryos at 120 and 200 min. White arrowheads indicate junctions enriched in DE-cadherin or myosin II. (B and C) DE-Cadherin (B) and myosin II (C) polarization ratios over time in dorsal and ventral cells of control embryos. The DE-cadherin ratio is the mean intensity at edges perpendicular to the DV axis divided by that at edges parallel to it; the myosin II ratio is the inverse (STAR Methods). (D and E) DE-Cadherin (D) and myosin II (E) polarization ratios at 120 and 200 min in control dorsal, control ventral and *Dia<sup>CA</sup>* ventral cells. The red dashed line indicates a ratio of 1. (F) *pnr*-Gal4 > UAS-palmmKate (magenta) with DE-Cadherin::GFP (green). (F') Mean intensity of palmmKate per cell in each row (n=598 cells from one embryo). (G) DE-Cadherin::Tomato in *Dia<sup>CA</sup>* embryo at 200 min showing landmarks P1 (leading edge), P2 (third sensillum) and P3 (forth sensillum). Orange, blue and gray lines represent respectively dorsal domain (P1-P2 distance), ventral domain (P2-P3 distance) and total epidermis extension (P1-P3 distance). (G'') Mean lengths of the dorsal domain (P1-P2), ventral domain (P2-P3), and total epidermis extension (P1-P3) over time in control (dashed, N=5) and *Dia<sup>CA</sup>* (solid, N=3) embryos. (H) Segmented cells color-coded by cell width in control and *Dia<sup>CA</sup>* embryos at 120 and 200 min. The white line marks the dorsal-ventral boundary. (I) Cell width per cell row at 120 and 200 min in control and *Dia<sup>CA</sup>* embryos.

(B-C) Polarizations of DE-Cadherin (B) and myosin II (C) are presented as mean ± SEM (n =4353, N=6). (D-E) Boxes show the median and interquartile range of DE-Cadherin (D) and myosin II polarization ratios (controls, n=1508 N=6; *Dia<sup>CA</sup>*, n=549, N=3). Statistical significance was assessed using two-way ANOVA followed by Benjamini-Hochberg corrected Welch's t-tests (\*\*\* p < 0.001, ns). (I) Cell width data are presented as mean ± SEM (control, n=3399, N=4; *Dia<sup>CA</sup>*, n=2403, N=3). Scale bars: (A) 5 μm; (F, H) 10 μm.

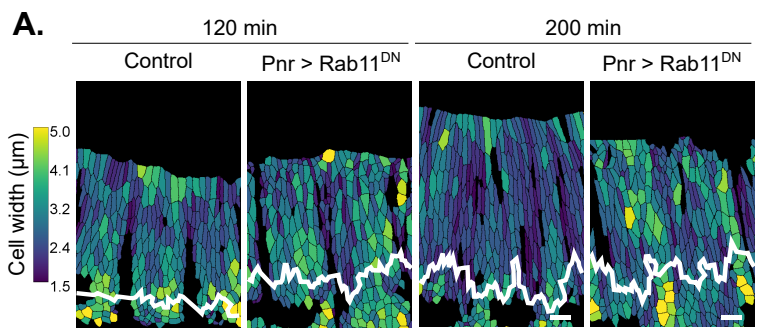

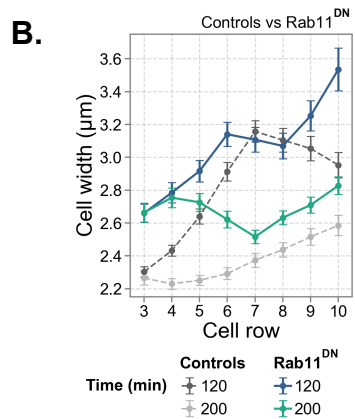

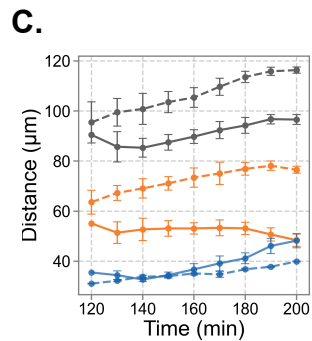

**Figure S3. Cell width and tissue deformation in Rab11<sup>DN</sup> embryos, related to Figure 3**

(A) Segmented cells color-coded by cell width in control and *pnr*-Gal4 > UAS-Rab11<sup>DN</sup> embryos at 120 and 200 min. The white line marks the dorsal–ventral boundary. (B) Cell width per row at 120 and 200 min in control (gray, dashed) and Rab11<sup>DN</sup> (colored) embryos. (C) Mean lengths of the dorsal domain (P1–P2), ventral domain (P2–P3), and total epidermis extension (P1–P3) over time in control (dashed, N=3) and Rab11<sup>DN</sup> (solid, N=3) embryos.

Maximal extension of P1–P3 was used to align embryos in time (STAR Methods). (B) Cell width data are presented as mean ± SEM (control, n=2791, N=3; Rab11<sup>DN</sup>, n=2867, N=3). Scale bars: (A) 10 μm.

#### Cell mechanics

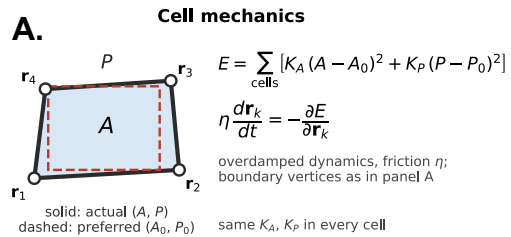

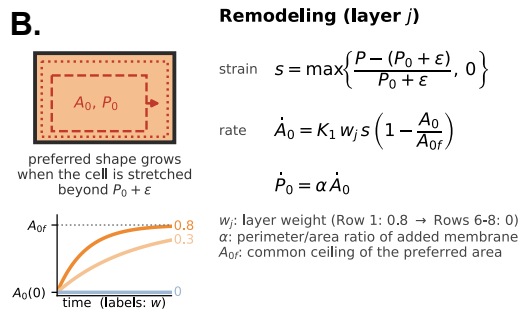

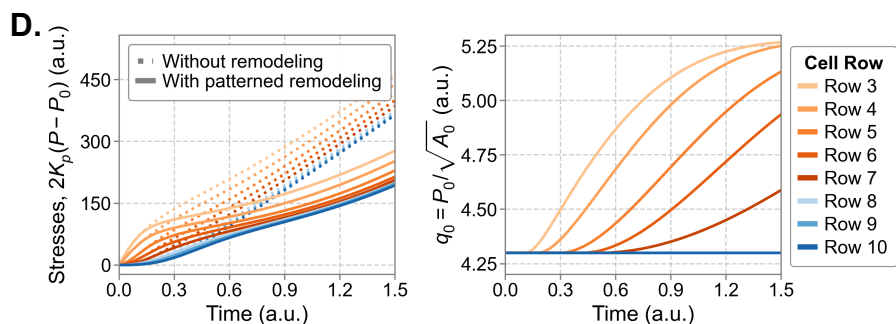

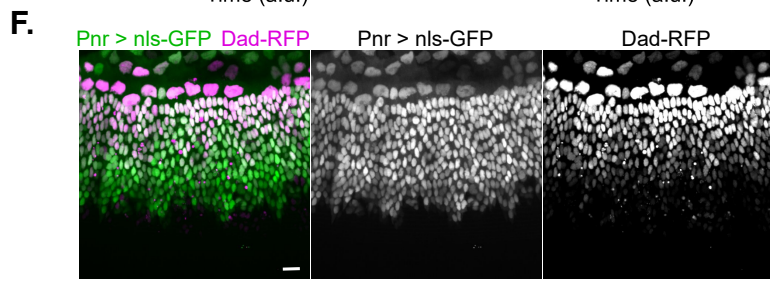

#### Without remodeling

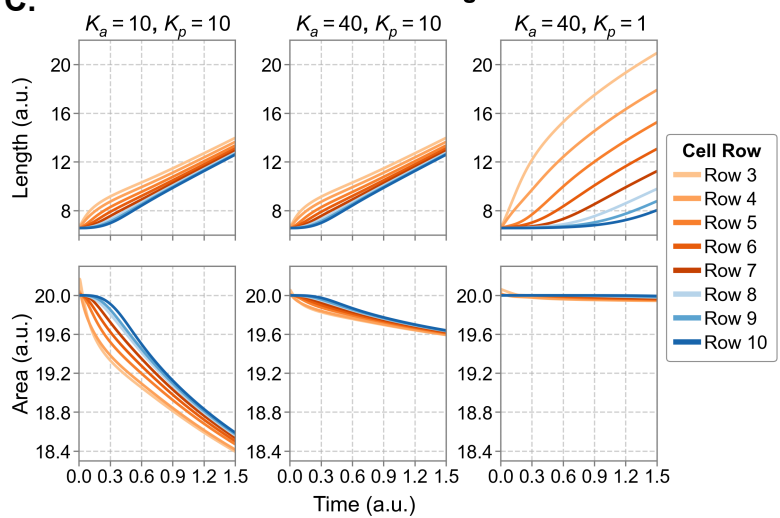

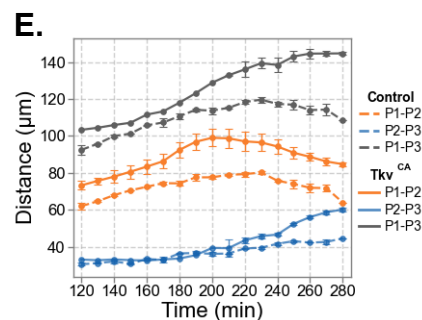

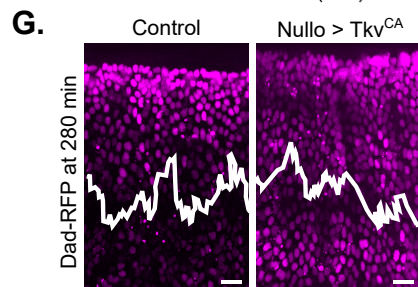

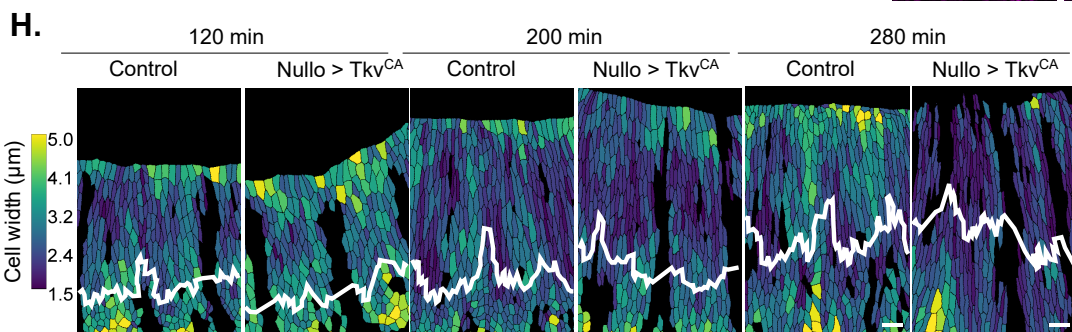

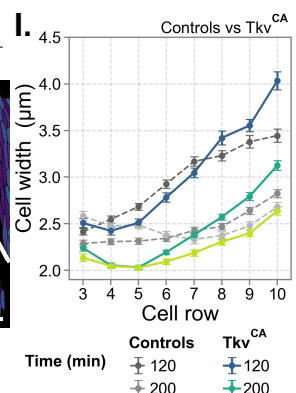

##### Figure S4. Vertex model and $Tkv^{CA}$ embryos, related to Figure 4

(A) Cell mechanics in the vertex model. Each cell has an actual area $A$ and perimeter $P$ (solid outline) and preferred values $A_0$ and $P_0$ (dashed outline). The energy and the overdamped equations of motion are indicated; $K_A$ and $K_P$ are identical for all cells. (B) Remodeling rule. When the perimeter of a cell exceeds $P_0 + \epsilon$, its preferred area grows at a rate proportional to the strain $s$ and to the row weight $w$, saturating at $A_{of}$; the preferred perimeter grows in proportion ($\alpha$). Bottom: $A_0$ over time for different weights. (C) Cell length and apical area over time for each row from the vertex model without remodeling, for $(K_A, K_P) = (10, 10)$, $(40, 10)$ and $(40, 1)$. Perimeter tension, $2K_P(P - P_0)$, over time for each row without remodeling (solid lines) and with patterned remodeling (dotted lines), using identical mechanical parameters ($K_A = 40$, $K_P = 10$). Orange, remodeling rows (3–7); blue, non-remodeling rows (8–10). (D) Target shape index $q_0 = P_0/\sqrt{A_0}$ over time for each row with patterned remodeling. Rows 8–10 do not remodel and overlap at $q_0 = 4.3$. (E) Mean lengths of the dorsal domain (P1–P2), ventral domain (P2–P3), and total epidermis extension (P1–P3) over time in control (dashed, $N=3$) and $Tkv^{CA}$ (solid, $N=3$) embryos. (F) Dad-RFP (magenta) and merge with *pnr*-Gal4 > UAS-nls-GFP (green). (G) Dad-RFP (magenta) in control and *nullo*-Gal4 > UAS-$Tkv^{CA}$ embryos at 280 min. The white line marks the dorsal–ventral boundary. Note that in $Tkv^{CA}$ embryos ventral cells show high Dad-RFP intensity. (H) Segmented cells color-coded by cell width in control and $Tkv^{CA}$ embryos at 120, 200 and 280 min. The white line marks the dorsal–ventral boundary. (I) Cell width per row in control and $Tkv^{CA}$ embryos.

Model quantities are in arbitrary units. (I) Cell width data are presented as mean ± SEM (control, $n = 4945$, $N=3$; $Tkv^{CA}$, $n = 4388$, $N=3$). Scale bars: (F-H) 10 μm.

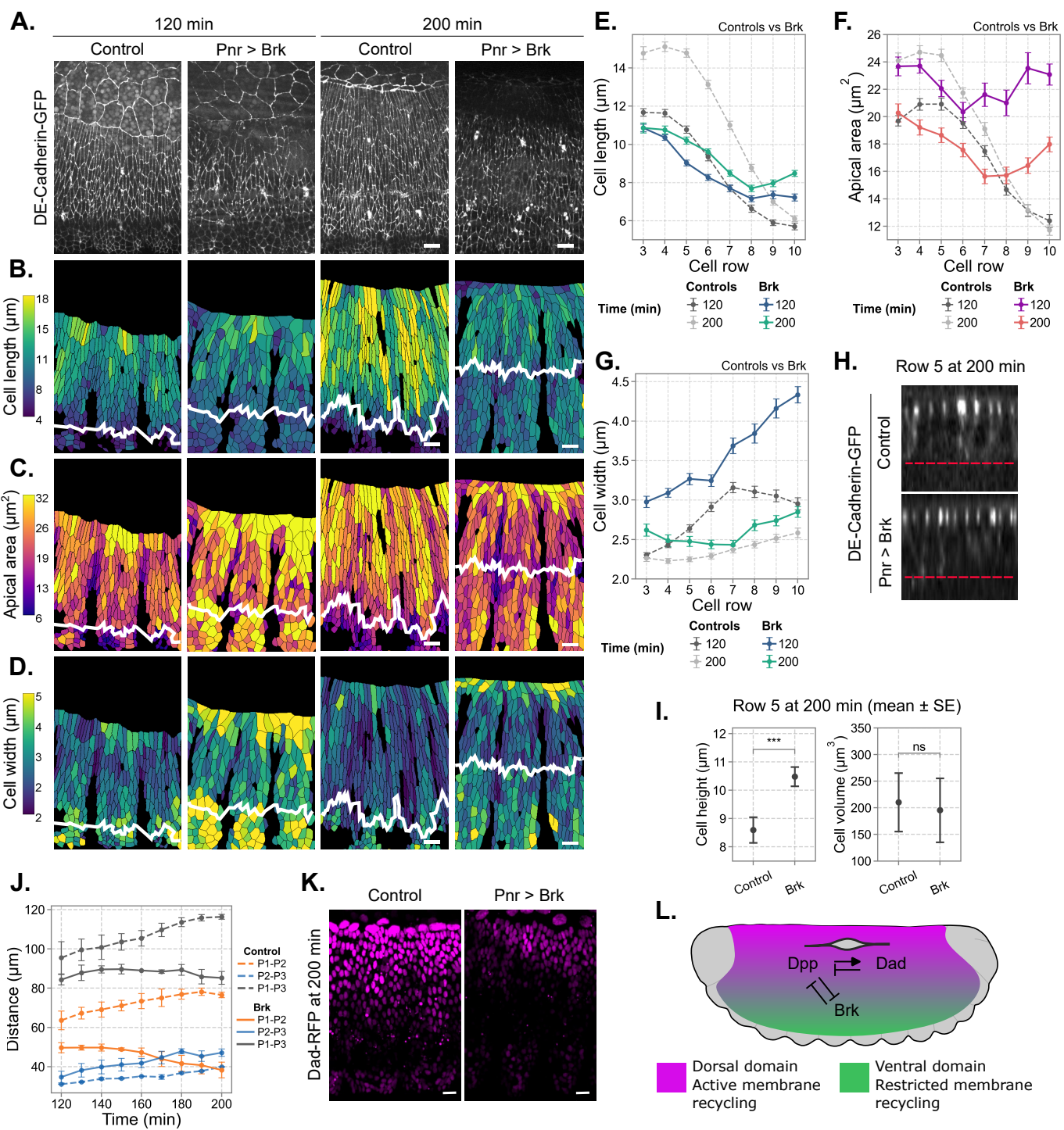

**Figure S5. Brinker expression in the dorsal epidermis prevents its elongation, related to Figure 4**

(A) Maximum-intensity projections of DE-Cadherin::GFP in control and *pnr-Gal4 > UAS-brk* embryos at 120 and 200 min. (B–D) Segmented cells color-coded by cell length (B), apical area (C) and cell width (D). The white line marks the dorsal–ventral boundary. (E–G) Cell length (E), apical area (F) and cell width (G) per row at 120 and 200 min in control (gray, dashed) and Brk (colored) embryos. (H) Orthogonal sections of row 5 at 200 min in control and Brk embryos. The red dashed line indicates the basal surface. (I) Cell height and volume of cells from row 5 at 200 min. (J) Mean lengths of the dorsal domain (P1–P2), ventral domain (P2–P3), and total epidermis extension (P1–P3) over time in control (dashed, N=3) and Brk (solid, N=3) embryos. (K) Dad-RFP (magenta) in control and *pnr-Gal4 > UAS-Brk* embryos at 200 min. Note that in Brk embryos the Dad-RFP intensity is lower. (L) Scheme of Brk–Dpp–Dad axis interaction in the epidermis. Dpp signaling in the dorsal domain (magenta) promotes active cell remodeling, whereas Brk signaling in the ventral domain (green) restricts cell remodeling by reducing Dpp activity.

Embryos were aligned on maximal extension of the epidermis ($t = 200$ min; STAR Methods). (E–G) Apical area, cell length and cell width data are presented as mean ± SEM (control, $n = 2791$, $N=3$; Brk, $n = 1727$, $N=3$). (I) Cell height data are presented as mean ± SEM (control, $n=11$, $N=3$; Brk, $n=10$, $N=3$). Cell volume was estimated as the product of mean cell height and mean apical area. Statistical comparisons of cell height and volume were performed using a z-test based on group means and standard errors (height, \*\*\*$p < 0.001$; volume, ns). Scale bars: (A–C, K) $10\ \mu\text{m}$; (H) $5\ \mu\text{m}$.

### Description of Additional Supplementary Files

**Movie S1.** Time-lapse of a control embryo during the whole dorsal closure process. DE-cadherin is labelled with Tomato so that we can observe apical cell shape and tissue deformation. Note the progressive and patterned elongation and increase in apical area of the dorsal epidermal cells. Scale bar 20 $\mu\text{m}$.

**Movie S2.** Time-lapse of a control embryo during the whole dorsal closure process. Myosin II is labelled with GFP allowing us to observe its polarization during tissue deformation. The embryo shown here is the same as in Movie S1. Scale bar 20 $\mu\text{m}$.

**Movie S3.** Time-lapse of an embryo expressing $\text{Dia}^{\text{CA}}$ in the *pannier* domain (dorsal cells). DE-cadherin is labelled with Tomato so that we can observe apical cell shape and tissue deformation. $\text{Dia}^{\text{CA}}$ expression in the dorsal domain blocks dorsal cell elongation and apical area increase. Consequently, the ventral cells elongate more than in control embryos, but with a similar apical area. At 240 min, note the severe defect in dorsal closure with the epidermis detaching from the amnioserosa, resulting in the embryo opened phenotype. Scale bar 20 $\mu\text{m}$.

**Movie S4.** Time-lapse of an embryo expressing $\text{Dia}^{\text{CA}}$ in the *pannier* domain (dorsal cells). Myosin II is labelled with GFP so that we can observe its polarization. In the absence of elongation of the dorsal cells and with increased elongation of the ventral cells (see Movie S3), the ventral cells show high Myosin II polarization upon greater stress, comparable to that of the dorsal cells in control embryos. The embryo shown is the same as in Movie S3. Scale bar 20 $\mu\text{m}$.

**Movie S5.** Time-lapse of an embryo expressing $\text{Rab11}^{\text{DN}}$ in the *pannier* domain (dorsal cells). DE-cadherin is labelled with GFP so that we can observe apical cell shape and tissue deformation. $\text{Rab11}^{\text{DN}}$ expression in the dorsal domain blocks dorsal cell elongation and apical area increase. Scale bar 20 $\mu\text{m}$.

**Movie S6.** Time-lapse of an embryo expressing $\text{Tk}^{\text{CA}}$ in the *nullo* domain (whole epidermis). DE-cadherin is labelled with GFP so that we can observe apical cell shape and tissue deformation. Homogenization of Dpp signaling in the epidermis through $\text{Tk}^{\text{CA}}$ expression homogenizes cell shapes across the embryo DV axis. Scale bar 20 $\mu\text{m}$.

**Movie S7.** Time-lapse of an embryo expressing Brk in the *pannier* domain (dorsal cells). DE-cadherin is labelled with GFP so that we can observe apical cell shape and tissue deformation. Brk expression in the dorsal cells impairs their elongation. Scale bar 20 $\mu\text{m}$.
